# The ADP-heptose biosynthesis enzyme GmhB is a conserved Gram-negative bacteremia fitness factor

**DOI:** 10.1101/2022.03.08.483568

**Authors:** Caitlyn L. Holmes, Sara N. Smith, Stephen J. Gurczynski, Geoffrey B. Severin, Lavinia V. Unverdorben, Jay Vornhagen, Harry L. T. Mobley, Michael A. Bachman

## Abstract

*Klebsiella pneumoniae* is a leading cause of Gram-negative bacteremia, which is a major source of morbidity and mortality worldwide. Gram-negative bacteremia requires three major steps: primary site infection, dissemination to the blood, and bloodstream survival. Since *K. pneumoniae* is a leading cause of healthcare-associated pneumonia, the lung is a common primary infection site leading to secondary bacteremia. *K. pneumoniae* factors essential for lung fitness have been characterized, but those required for subsequent bloodstream infection are unclear. To identify *K. pneumoniae* genes associated with dissemination and bloodstream survival, we performed insertion site sequencing (InSeq) using a pool of >25,000 transposon mutants in a murine model of bacteremic pneumonia. This analysis revealed the gene *gmhB* as important for either dissemination from the lung or bloodstream survival. In *Escherichia coli*, GmhB is a partially redundant enzyme in the synthesis of ADP-heptose for the lipopolysaccharide (LPS) core. To characterize its function in *K. pneumoniae*, an isogenic knockout strain (Δ*gmhB*) and complemented mutant were generated. During pneumonia, GmhB did not contribute to lung fitness and did not alter normal immune responses. However, GmhB enhanced bloodstream survival in a manner independent of serum susceptibility, specifically conveying resistance to spleen-mediated killing. In a tail-vein injection of murine bacteremia, GmhB was also required by *K. pneumoniae*, *E. coli* and *Citrobacter freundii* for optimal bloodstream survival. Together, this study identifies GmhB as a conserved Gram-negative bacteremia fitness factor that acts through LPS-mediated mechanisms to enhance bloodstream survival.

**IMPORTANCE:** *Klebsiella pneumoniae* frequently causes healthcare-associated infections including pneumonia and bacteremia. This is particularly concerning due to emerging antimicrobial resistance and the propensity for bacteremia to initiate sepsis, which has high mortality and is the most expensive hospital-treated condition. Defining mechanisms of bloodstream survival is critical to understanding the pathology of bacteremia and identifying novel targets for future therapies. In this study, we identified the *K. pneumoniae* enzyme GmhB as a bloodstream-specific fitness factor that enables the bacteria to survive in the spleen but is dispensable in the lung. Furthermore, GmhB is also needed by the related bacterial pathogens *Escherichia coli* and *Citrobacter freundii* to cause bacteremia. Conserved bacteremia fitness factors such a GmhB could be the basis for future therapeutics that would alleviate significant disease caused by from multiple diverse pathogens.

## INTRODUCTION

Gram-negative bacteremia is a significant cause of global morbidity and mortality largely due to progression to sepsis, defined as life threatening organ dysfunction resulting from a dysregulated host response to infection (1). Gram-negative pathogens underlie 43% of clinical bloodstream infections with a small number of species, including *Escherichia coli*, *Klebsiella pneumoniae, Citrobacter freundii*, and *Serratia marcescens*, contributing to the majority of cases (2, 3). Of these species, *K. pneumoniae* is the second most common species causing Gram-negative bacteremia and the third most prevalent cause of all bloodstream infections (2). Although *K. pneumoniae* can be a commensal species (4, 5), it is also an opportunistic pathogen. This is especially relevant in healthcare-associated infections where *K. pneumoniae* is a leading source of disease (6). The Centers for Disease Control and Prevention have repeatedly classified carbapenem-resistant *Enterobacterales*, including *K. pneumoniae*, as an urgent public health threat due to antibiotic resistance (7, 8). Bacteremia from antibiotic-resistant *K. pneumoniae* can be extremely difficult to treat and is associated with a high mortality rate.

The pathogenesis of Gram-negative bacteremia involves three main phases: primary site infection, dissemination, and bloodstream survival (3). First, bacteria must invade primary sites of infection or colonization and evade local host responses. Second, pathogens disseminate across host barriers to gain bloodstream access, a process that varies based on the initial site. Navigation across barriers may include strategies to invade or disrupt site-specific epithelial cells, endothelial cells, and cellular junctions. Third, bacteria must exercise metabolic flexibility and resist host defenses in the bloodstream to adapt in a new environment. In circulation, bacteria passage through blood filtering organs, like the spleen and liver, which may act as additional sites of infection from which dissemination can occur. Defects at initial sites do not always predict fitness at secondary sites (9, 10), and apparent lack of fitness at secondary sites may be confounded by defects at the initial site. Therefore, observed overlap between primary site and bloodstream fitness genes highlight the necessity to probe phases of bacteremia separately to correctly define stages relevant to pathogenesis (3). By carefully defining the bacterial factors required for each phase of bacteremia, we may identify therapeutic targets for interventions that prevent progression to bacteremia or treat it more effectively once it has occurred.

*K. pneumoniae* bacteremia is often secondary to pneumonia (6) and fitness factors for primary site infection in the lung have been extensively investigated. Capsular polysaccharide, siderophores, and synthesis of branched chain amino acids (11–13) are required for lung fitness. Additionally, the citrate (Si)-synthase GltA, and the acetyltransferase Atf3, are required (9, 10), highlighting the broad range of factors contributing to lung initial site fitness. Some fitness factors in the lung are also likely to be important in the bloodstream. Capsular polysaccharide is required to resist human serum complement, and siderophores are important for both dissemination from the lung and growth in human serum (12). However, factors that act specifically at the stages of dissemination and bloodstream survival are unclear. Genes necessary for serum resistance have been described *in vitro* and include cell wall integrity proteins, and multiple metabolic pathways (14, 15), but factors that resist host responses during bacteremia and allow growth within blood-filtering organs is unknown.

In the bloodstream, cell surface structures can defend bacteria from environmental threats like formation of the membrane attack complex or antimicrobial peptides. Of these, lipopolysaccharide (LPS) is a defining cellular envelope structure of Gram-negative species that governs many environmental interactions and aids in resistance to stress. Major components of the LPS molecule include O-antigen, outer core, inner core, and lipid A. LPS alterations can increase vulnerability to environmental threats (16), and inner core mutations can enhance susceptibility to hydrophobic agents (16–18). Since LPS can also interact with host Toll-like receptor 4 to initiate innate immune responses, it is likely that *K. pneumoniae* LPS plays a complex role in host-pathogen interactions during bacteremia.

To identify factors required for lung dissemination and bloodstream survival, we used transposon insertion site sequencing (InSeq) in a murine model of bacteremic pneumonia. We identified and validated the LPS core biosynthesis gene *gmhB* as involved in dissemination and bloodstream survival, the two late phases of bacteremia, but dispensable for initial site fitness in the lung. We also showed that GmhB is a conserved bloodstream survival factor across multiple Gram-negative pathogens.

## RESULTS

### Transposon insertion site sequencing identifies *K. pneumoniae* GmhB as a bacteremia fitness factor

To identify *K. pneumoniae* factors influencing dissemination and bloodstream survival, InSeq was used to detect genes associated with fitness defects in the spleen but dispensable for lung fitness. The *K. pneumoniae* strain KPPR1 causes bacteremic pneumonia in a well-established murine model (13, 19). In a previous study to identify interactions between *Klebsiella* and the innate immune protein Lipocalin 2 during pneumonia (10), we used a KPPR1 transposon library representing ~25,000 unique insertions with ~99% genome coverage to infect *Lcn2*^+/+^ and *Lcn2*^-/-^ mice (11, 20). Here, we evaluated the dissemination of mutants to the spleen at 24 hours from the same experiment. We noted that the *Lcn2*^-/-^ mice had greater dissemination to the spleen than *Lcn2*^+/+^ mice (Supplementary Figure 1), suggesting a wider bottleneck in dissemination from the lung that could enable higher recovery of transposon mutants in the spleen. Therefore, only *Lcn2*^-/-^ spleens were analyzed further (21).

To identify potential lung dissemination and bloodstream survival factors, we devised a stepwise approach to use the InSeq data from the spleens of *Lcn2*^-/-^ mice and eliminated genes with fitness defects in the lung or interactions with Lipocalin 2: Genes containing transposon insertions were compared between the inoculum, *Lcn2*^+/+^ lung, *Lcn2*^-/-^ lung, and *Lcn2*^-/-^ spleen output pools. Of the 3,707 mutated genes shared across the input and each output pool, 1,489 contained four or more unique transposon insertions (*i.e*., median number of unique insertions per gene) and were used for subsequent selection steps. To eliminate genes influencing lung fitness, transposon mutants with similar abundance (q>0.05) between the inoculum and *Lcn2*^+/+^ mouse lungs were retained. To eliminate genes that interact with Lipocalin 2 in the lungs, only transposon mutants with similar recovery (q>0.05) between the *Lcn2*^+/+^ and *Lcn2*^-/-^ lung output pools were retained. To identify factors involved in either the phase of lung egress or bloodstream survival, transposon mutants were selected with a significant difference in abundance between the *Lcn2*^-/-^ lung and *Lcn2*^-/-^ spleen output pools (q<0.05). This InSeq selection process resulted in 18 genes with transposon insertions (Supplemental Table 1) as candidates for encoding dissemination and bloodstream survival factors.

Six genes with a high ratio in read difference between the lung and spleen were chosen for validation. Isogenic knockouts of the open reading frames of *VK055_4727*, *VK055_2040*, *VK055_4483*, *ulaA*, *gmhB*, and *prlC* were generated by Lambda Red mutagenesis (22). None of the encoded factors were required for *K. pneumoniae in vitro* replication or fitness, as knockouts had growth rates similar to those of wild-type KPPR1 in rich LB and minimal (M9+Glucose) media (Supplementary Figure 2A, C). Additionally, each knockout was able to compete *in vitro* against wild-type KPPR1 with no apparent defects in both media conditions (Supplementary Figure 2B, D). To validate the defect of each mutant in causing bacteremia, 1:1 coinfections of KPPR1 against each isogenic knockout in a bacteremic pneumonia model were performed with *Lcn2*^-/-^ mice. Competitive indices were calculated 24 hours post inoculation based on bacterial burden of each strain (Supplemental Figures 3–4). The Δ*gmhB* mutant had a slight fitness defect in the lung with a significantly greater defect in the spleen (Supplementary Figure 3E). This significant difference in fitness between sites indicates that GmhB is important for lung dissemination, bloodstream survival, or both steps of bacteremia. In contrast, VK055_4727, VK055_2040, VK055_4483, and UlaA were dispensable for bacteremia at all phases (Supplementary Figure 3A-D). PrlC contributes to initial site fitness (Supplementary Figure 3F), which may explain the similar fitness defect observed in the spleen.

### Multiple models of murine bacteremia support that GmhB enhances bloodstream fitness

Since Lcn2 can prevent *K. pneumoniae* pulmonary vasculature invasion (23), we confirmed that GmhB was required for dissemination and bloodstream survival in wild-type mice. Consistent with the InSeq data, the *gmhB* mutant had no fitness defect in the lungs of *Lcn2*^+/+^ mice after coinfections with KPPR1 and Δ*gmhB* (Figure 1A, Supplementary Figure 5A). In contrast, the *gmhB* mutant had a 24-fold mean fitness defect in the spleen and 104-fold defect in blood. Similar to co-infections, in independent infections the *gmhB* mutant had no defect in the lung but significant defects in the spleen and blood of infected mice (Figure 1B). To confirm that this fitness defect was attributable to disruption of *gmhB*, the mutant was complemented *in trans*. The empty plasmid vector had no effect on the results of competitive infections (Figure 1C). Plasmid carriage had slight effects on lung fitness, with Δ*gmhB* carrying the empty vector having slightly higher fitness, and Δ*gmhB* with the complementing plasmid having slightly lower fitness, in the lung (Figures 1C-D; Supplementary Figure 5B-C). In contrast, Δ*gmhB* with the empty vector was significantly defective for survival in the spleen and blood with plasmid derived *gmhB* complementation ameliorating this defect in the spleen and partially in the blood (Figure 1D). Combined, these results indicate that GmhB is necessary for lung dissemination, bloodstream survival, or both stages of bacteremia.

**Figure 1.**
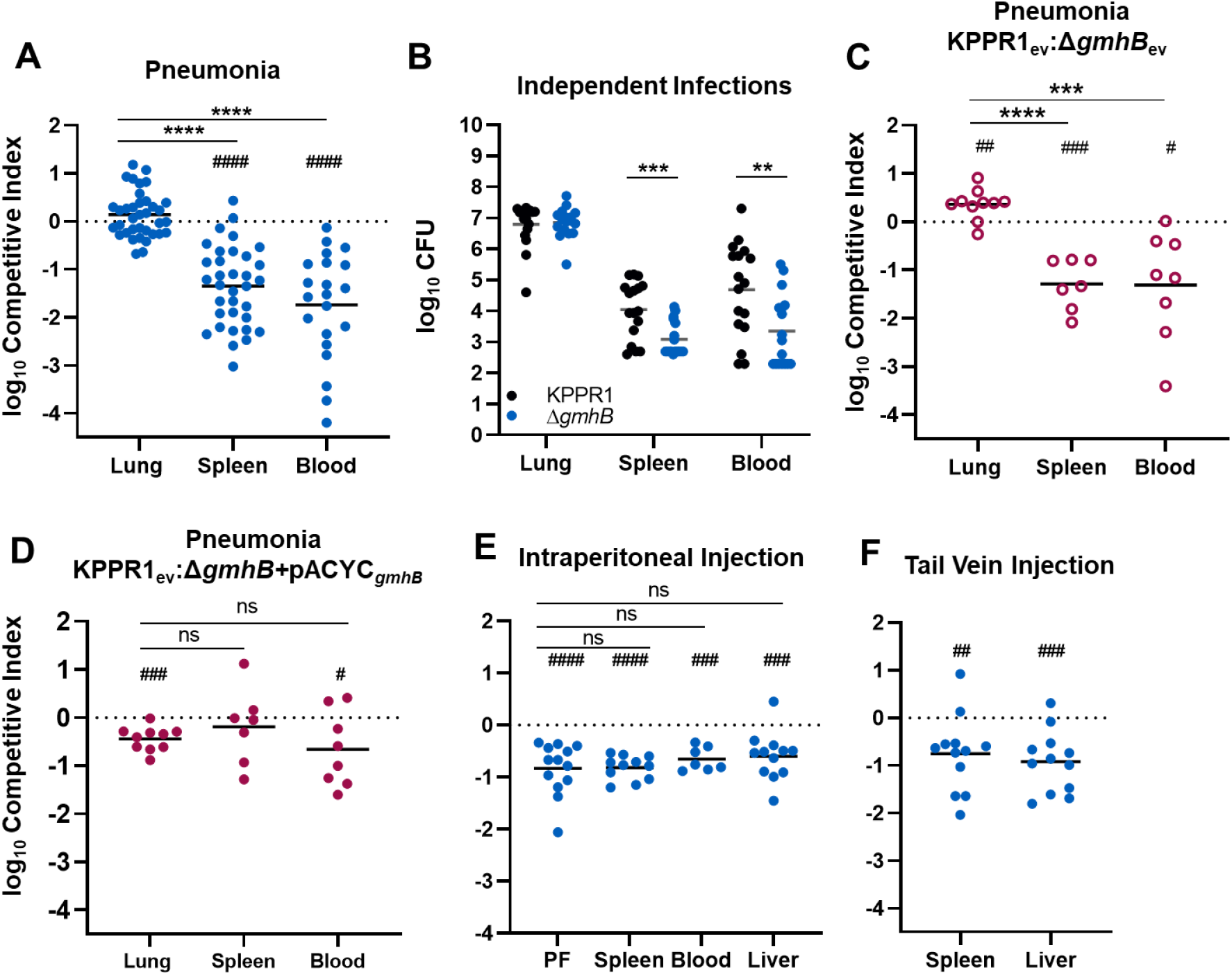
GmhB enhances lung dissemination and bloodstream survival. In a model of bacteremic pneumonia, mice were retropharyngeally inoculated with 1×10^6^ CFU *K. pnuemoniae* (A-D). To initiate dissemination from a lung-independent site, 1×10^3^ CFU was administered to the intraperitoneal cavity (E). For modeling direct bacteremia requiring no dissemination, 1×10^5^ CFU was administered via tail vein injection (F). The 1:1 inoculum consisted of KPPR1:Δ*gmhB* (A, E, F), KPPR1:Δ*gmhB* carrying empty pACYC vector (ev; C), or KPPR1_ev_:Δ*gmhB* with *gmhB* complementation provided on pACYC under the control of the native *gmhB* promoter (Δ*gmhB*+pACYC_*gmhB*_; D). Independent infections used either KPPR1 or Δ*gmhB* alone at a 1×10^6^ CFU dose (B). Mean log_10_ competitive index or CFU burden at 24 hours post infection is displayed. **p<0.01, ***p<0.001, ****p<0.0001 by unpaired t test; ^##^p<0.01, ^###^p<0.001, ^####^p<0.0001 by one sample t test with a hypothetical value of zero. For each group, n≥7 mice in at least two independent infections, PF=peritoneal fluid.

To determine if GmhB enhances dissemination from the lung specifically, a bacteremia model involving an independent initial site was used. A KPPR1 and Δ*gmhB* coinfection was performed by intraperitoneal injection and competitive indices were calculated after 24 hours (Figure 1E, Supplementary Figure 5D). Unlike the lung model, the *gmhB* mutant was defective in initial site fitness within the peritoneal cavity and a similar fitness defect was observed in the spleen, liver, and blood. Therefore, GmhB influences initial phase fitness in a site-specific manner. This initial site defect in the intraperitoneal model may mask defects in bloodstream survival. To measure fitness in the third phase of bacteremia, a tail vein injection model was used that bypasses the initial site and dissemination steps. Based on coinfections using a tail vein injection with competitive indices calculated after 24 hours (Figure 1F, Supplementary Figure 5E), the *gmhB* mutant had a significant fitness defect in both the spleen and liver. Considering the data across three distinct models of bacteremia, GmhB is consistently necessary for bloodstream survival. It is dispensable for initial site infection in the lung but important in the peritoneal cavity, suggesting site-specific fitness. The contribution of GmhB to bloodstream survival may explain the strong defect in dissemination observed in pneumonia model, but we cannot rule out a specific contribution for egress from the lung.

### GmhB does not modulate lung inflammation elicited by *K. pneumoniae* during pneumonia

GmhB is a D,D-heptose 1,7-bisphosphate phosphatase involved in biosynthesis of ADP-heptose (24–26), which is a structural component of the LPS core. ADP-heptose is synthesized through a five-part enzymatic cascade modifying the precursor sedoheptulose 7-phosphate. GmhB is the third enzyme in this reaction, serving to dephosphorylate D-glycero-β-D-manno-heptose 1,7-bisphosphate (HBP) to produce D-glycero-β-D-manno-heptose 1-monophosphate (HMP1) (25). Perhaps because LPS is a conserved virulence factor in Gram-negative bacteria, ADP-heptose is also a soluble pro-inflammatory mediator (27). Soluble ADP-heptose can be recognized by the host cytosolic receptor alpha kinase 1 (ALPK1) (27), resulting in the formation of TIFAsomes, upregulation of NF-κb signaling, and inflammatory influx (28–31). We have previously observed that lung inflammation contributes to dissemination of *K. pneumoniae* from the lung to the bloodstream (12, 23). If lung dissemination is GmhB-dependent, then perhaps *K. pneumoniae* relies on soluble ADP-heptose to induce an immune response during pneumonia that enables egress from the lungs.

To measure the contribution of GmhB to lung inflammation, KPPR1 and Δ*gmhB* were used in the murine pneumonia model and lung homogenates were surveyed for immune cell recruitment and cytokine activation associated with ADP-heptose signaling (31). As expected, neutrophils and monocytes were the most prominent cell types recruited to the lung during *K. pneumoniae* infection (Figure 2A, Supplementary Figure 6) (32–34). Monocytic-myeloid derived suppressor cells (M-MDSCs), which alter the lung immune environment during *K. pneumoniae* infection (35, 36), were present after infection, but not in a GmhB-dependent manner. Alveolar macrophages, eosinophils and dendritic cells were detected by flow cytometry but the abundance of these cell types was not altered by *K. pneumoniae* infection. Importantly, GmhB did not influence the overall CD45^+^ cell abundance in the lung during pneumonia, nor did GmhB alter the profile of any prominent immune cell subset after infection (Figure 2A). We also measured the abundance of TNFα, GM-CSF, RANTES, MCP-3, MIP-1α, and MIP-1β, which are associated with signaling via the ADP-heptose/ALPK1/NF-κB axis (31), in lung homogenates. Abundance of each analyte was increased after *K. pneumoniae* infection, yet GmhB did not influence signaling by this axis (Figure 2B). Therefore, inflammation during *K. pneumoniae* lung infection is not GmhB-dependent, as measured by immune cell recruitment and signaling through ADP-heptose/ALPK1/NF-κB associated cytokines. The influence of GmhB on dissemination and bloodstream survival is likely independent of lung inflammatory responses.

**Figure 2.**
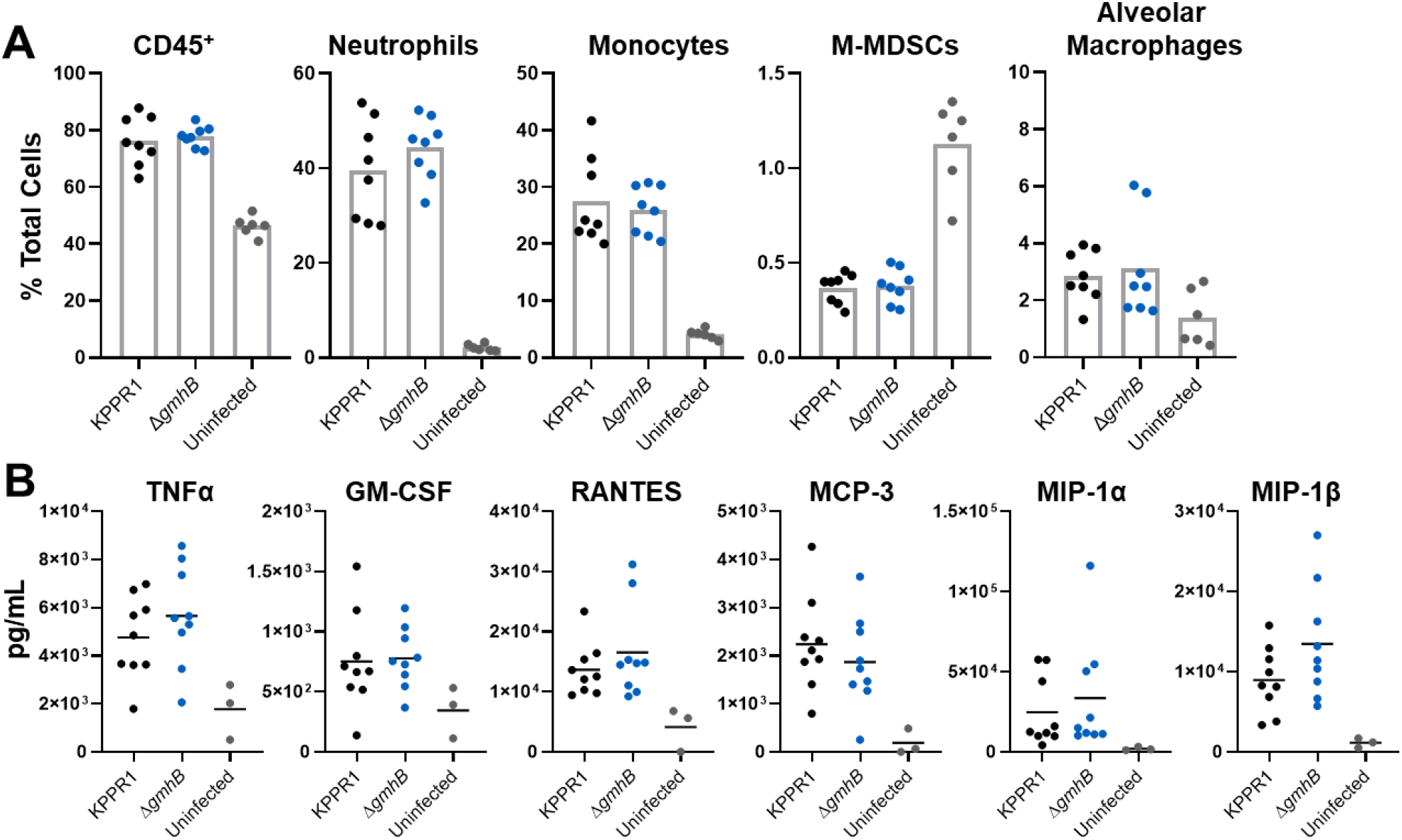
GmhB does not alter normal immune responses during *K. pneumoniae* lung infection. In a model of bacteremic pneumonia, mice were retropharyngeally inoculated with 1×10^6^ CFU of either KPPR1 or Δ*gmhB*. After 24 hours, lungs were prepared for flow cytometry using 1.5×10^6^ cells/lung. Comparisons between immune cell populations for KPPR1 or Δ*gmhB* infected or uninfected mice are displayed for relevant subsets (A). Cytokines associated with ADP-heptose/ALPK1 signaling were detected from lung homogenates using ELISA (B). For each infected group, n=8-9 mice, and for each uninfected group, n=3-6. Each panel represents infections from at least two independent experiments; no comparisons were significant by unpaired t test between KPPR1 and Δ*gmhB*.

### GmhB enhances bloodstream survival by mediating spleen fitness

Given that GmhB enhanced *K. pneumoniae* bloodstream survival during direct bacteremia (Figure 1F) and did not alter inflammation in the lungs (Figure 2), we investigated the direct role that it may play on bacterial fitness. Disruption of GmhB during ADP-heptose biosynthesis can influence LPS structure in *E. coli* (25, 26), and LPS core alterations may enhance serum susceptibility (24, 37). To determine if GmhB conveys resistance to serum killing, KPPR1 and Δ*gmhB* were exposed to active human and murine serum. An Δ*rfaH* acapsular mutant was used as a control that is highly susceptible to human serum killing (11). In contrast to RfaH, GmhB was dispensable for resistance to human serum-mediated killing (Figure 3A). Unlike human serum, murine serum was unable to elicit killing in any strain and may lack the ability to form an active membrane attack complex against *K. pneumoniae* (Figure 3B), a phenomenon observed in other Gram-negative species (38). Additionally, GmhB was not required for growth in active human serum (Figure 3C). To rule out subtle differences in fitness in human serum, competitive survival assays were performed in human serum. This also showed no defect of the *gmhB* mutant (Figure 3D, Supplementary Figure 7). Thus, the bloodstream survival advantage conveyed by GmhB is likely independent of the ability to resist complement-mediated killing or to replicate in serum.

**Figure 3.**
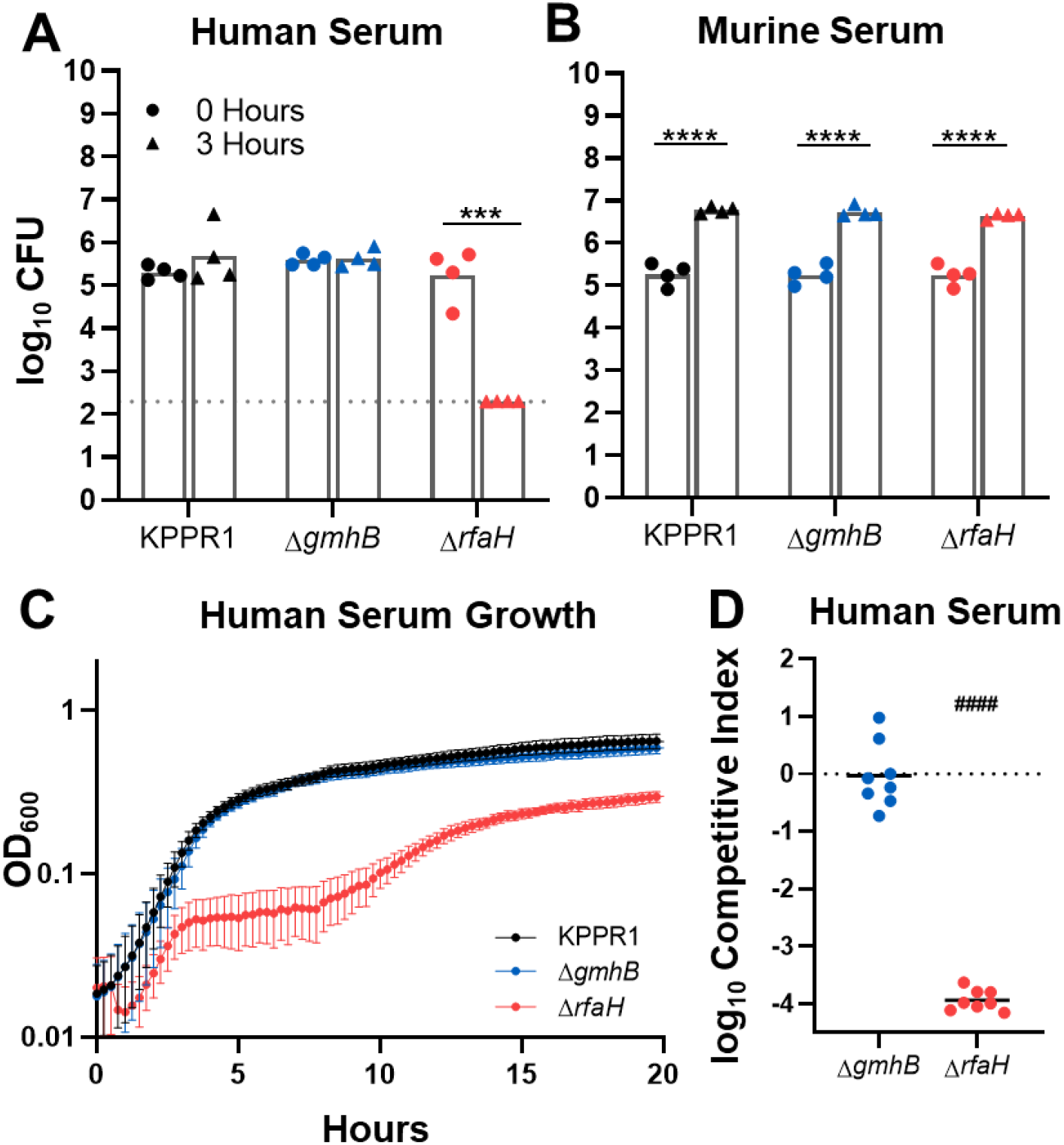
Bloodstream fitness conveyed by GmhB is serum independent. Serum susceptibility was compared after 3 hours for 1×10^5^ CFU KPPR1, Δ*gmhB*, and Δ*rfaH* in active human (A) or murine (B) serum. *K. pneumoniae* strains were grown in M9+20% active human serum and the OD_600_ was measured every 15 minutes for 20 hours (C). Competition assays were performed *in vitro* using active human serum (D) using a 1:1 mixture of 1×10^5^ KPPR1 and either Δ*gmhB* or Δ*rfaH*. Mean log_10_ competitive index compared to wild-type KPPR1 at 3 hours post infection is displayed. ***p<0.001, ****p<0.0001 by unpaired t test with n=4 (A-B) and limit of detection is represented by the dotted line. For D, p<0.0001 by one sample t test with a hypothetical value of zero and n=8.

During bacteremia, *Klebsiella* pass through blood filtering organs, like the liver and spleen, and GmhB conveyed a fitness advantage in these organs *in vivo* (Figure 1F). Since the fitness defects of Δ*gmhB* during bacteremia are not explained by fitness in serum, we performed *ex vivo* competition assays in uninfected murine spleen and liver homogenates. GmhB was necessary for complete fitness in spleen homogenate (Figure 4A, Supplementary Figure 7). Further, the magnitude of GmhB fitness loss in *ex vivo* spleen homogenate was similar to that observed *in vivo* using tail vein injections (Figure 1F). RfaH was dispensable for spleen homogenate fitness (Figure 4A) suggesting that capsule is not required for splenic survival. Furthermore, GmhB was dispensable for hypermucoviscosity (39) (Supplementary Figure 8). Despite finding a fitness defect and fewer Δ*gmhB* CFU in the liver during infection (Figures 1E,F and Supplementary Figure 5D, E), GmhB was dispensable for liver fitness *ex vivo* (Figure 4B). Similar to its neutral fitness in the lung, the *gmhB* mutant had no defect in lung homogenate *ex vivo* (Figure 4C). These data indicate that GmhB contributes to bacteremia fitness during the phase of bloodstream survival through spleen-specific interactions.

**Figure 4.**
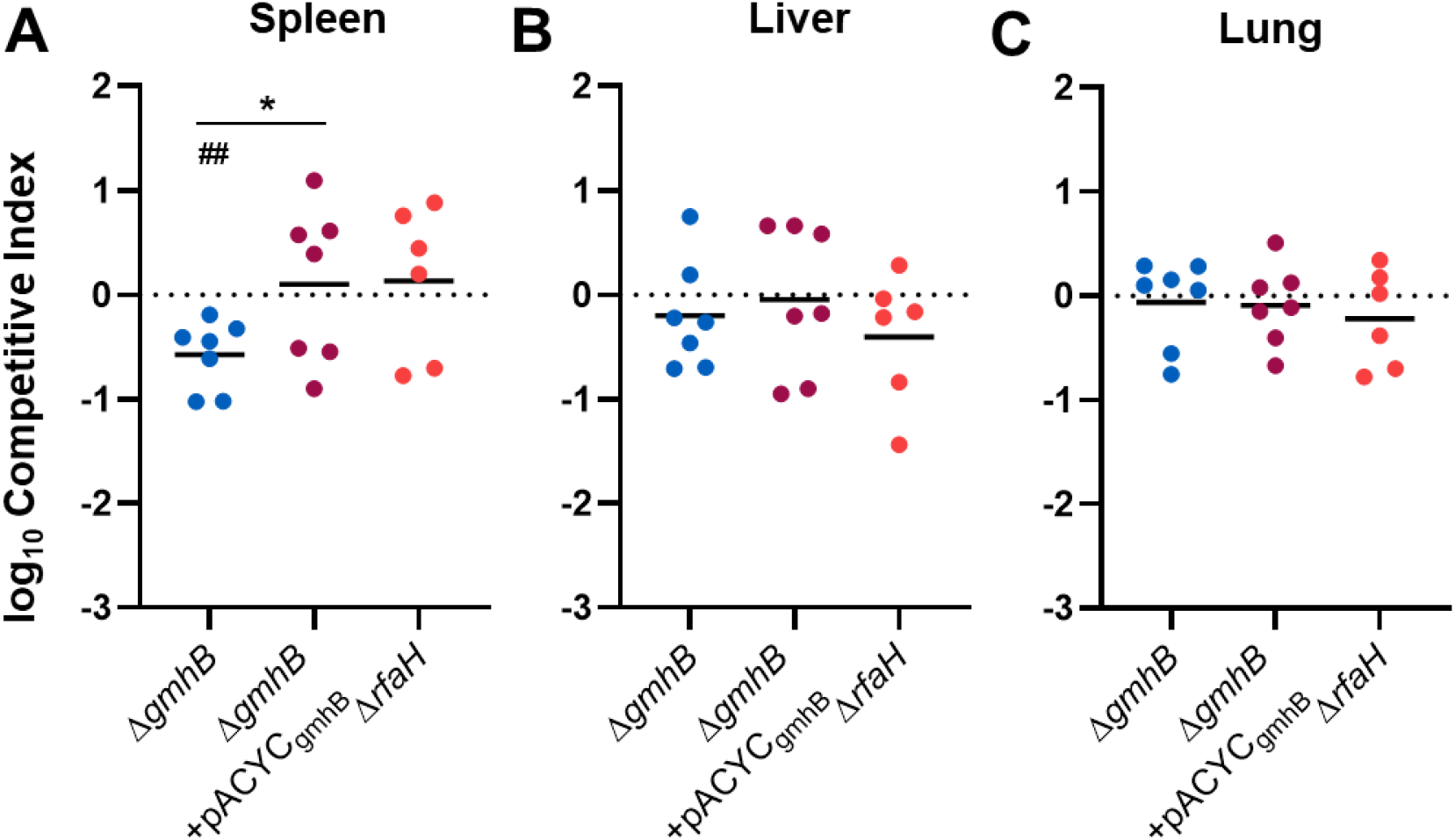
Bloodstream fitness conveyed by GmhB involves interactions in the spleen. Competition assays were performed *ex vivo* in murine spleen (A), liver (B), or lung (C) homogenate using a 1:1 mixture of 1×10^5^ KPPR1 and either Δ*gmhB*, Δ*gmhB*+pACYC_*gmhB*_, or Δ*rfaH*. Mean log_10_ competitive index compared to wild-type KPPR1 at 3 hours post inoculation is displayed. *p<0.05, by unpaired t test comparing KPPR1 and Δ*gmhB;* ^##^<0.01, by one sample t test with a hypothetical value of zero and n=6-7.

### GmhB is required for normal *K. pneumoniae* LPS composition

GmhB contributes to LPS structure through synthesis of ADP-heptose, a major component of the inner core region. In *E. coli*, GmhB is required for normal LPS composition; GmhB-deficient strains produce a mixed phenotype of full length and stunted LPS molecules (26). This partial defect is attributed to an uncharacterized enzyme that is partially redundant for GmhB function. In other species, disruption of ADP-heptose integration into LPS results in stunted molecules with minimal O-antigen (17, 18). To determine the impact of *gmhB* deletion on *K. pneumoniae* surface structure, LPS from KPPR1, Δ*gmhB*, and Δ*gmhB*+pACYC_*gmhB*_ was isolated and analyzed using electrophoresis. Wild-type KPPR1 LPS produces prominent O-antigen laddering patterns similar to the pattern of the *E. coli* LPS standard (Figure 5). The *K. pneumoniae* strain Δ*galU* (39, 40) lacks prominent O-antigen and can be used to identify regions corresponding to core polysaccharides. In three prominent core banding regions, differences were observed between wild-type KPPR1 and Δ*gmhB*. Specifically, there was decreased band intensity in heavier bands (Regions A and B) and the appearance of banding in Region C. These changes were reversed upon *gmhB* complementation. This result indicates that GmhB is required for normal *K. pneumoniae* LPS structure. Similar to *E. coli*, GmhB is not absolutely required for LPS synthesis as O-antigen laddering is still detected even in the absence of this enzyme.

**Figure 5.**
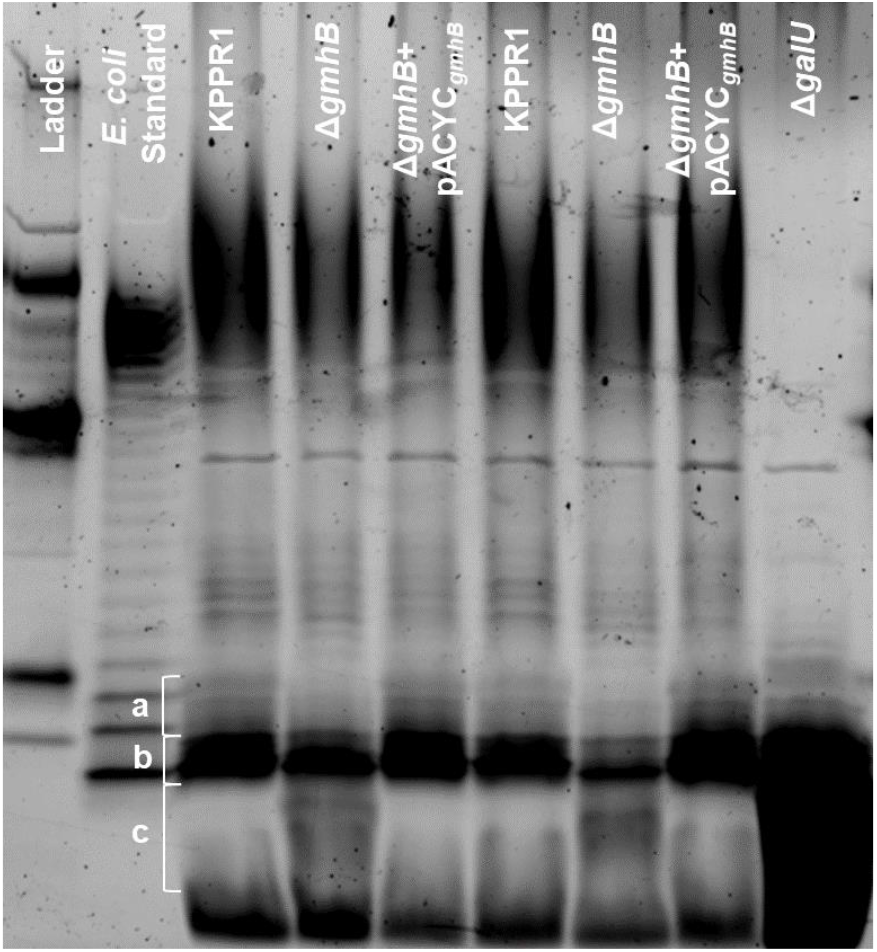
GmhB is required for normal LPS composition. LPS from 1×10^9^ CFU of KPPR1, Δ*gmhB*, Δ*gmhB*+pACYC_*gmhB*_, or Δ*galU* was isolated and 10μL of yield was analyzed by polyacrylamide electrophoresis. LPS core regions in interest are labeled in a, b, and c. The gel displayed is representative of three independent trials, duplicate lanes represent independent LPS preparations. The CandyCane glycoprotein molecular weight standard is displayed in the left lane.

### GmhB is a conserved bloodstream fitness factor across multiple clinically relevant Gram-negative bacteremia pathogens

GmhB is highly conserved across *Enterobacterales*, which compose the majority of Gram-negative bacteremia pathogens. To address the requirement of GmhB in bloodstream fitness across multiple species, tail vein injections were performed using a coinfection of wild type *E. coli* CFT073 or *C. freundii* UMH14 and corresponding *gmhB* mutants *CFT073:tn::gmhB* (42) and UMH14Δ*gmhB*, respectively. GmhB was required for bloodstream survival in both *E. coli* and *C. freundii* as measured in the spleen and liver (Figure 6, Supplementary Figure 9). Additionally, GmhB is a predicted essential gene for *S. marcescens* survival (43). These results reveal that GmhB is a conserved bloodstream fitness factor across multiple clinically relevant Gram-negative bacteremia pathogens.

**Figure 6.**
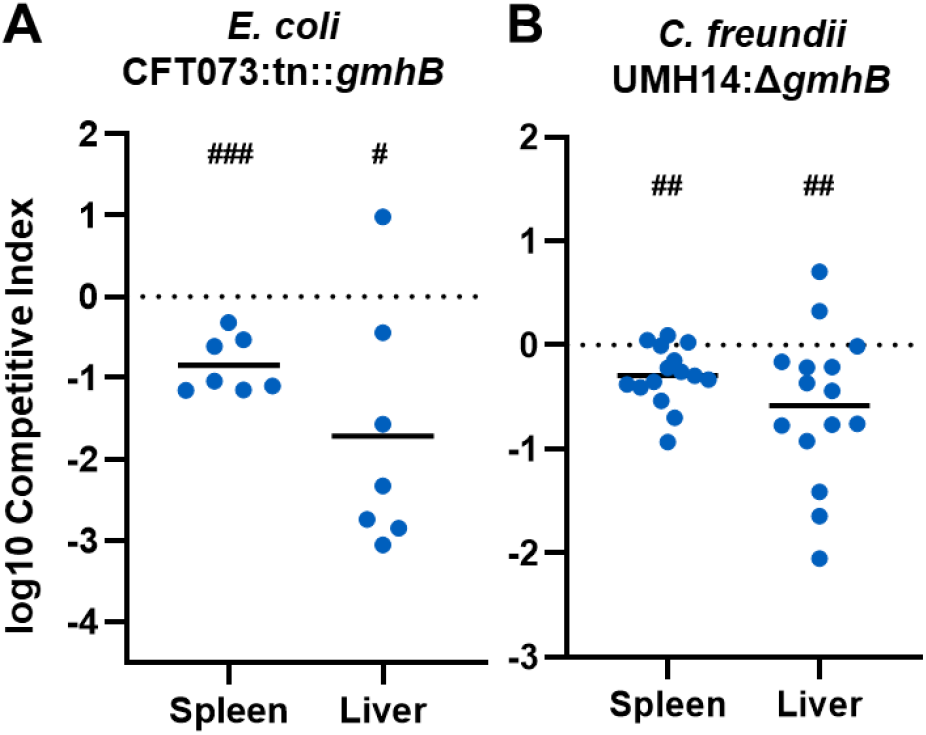
GmhB is required for bloodstream fitness across multiple Gram-negative species. In a model of bacteremia, 1×10^7^ CFU of *E. coli* CFT073 (A) or 7.5×10^7^ CFU *C. freundii* UMH14 (B) was administered via tail vein injection. The 1:1 inoculum consisted of CFT073:tn::*gmhB* (A) or 1:2 inoculum of UMH14:Δ*gmhB* (B). Mean log_10_ competitive index or CFU burden at 24 hours post infection is displayed. ^#^p<0.05, ^##^p<0.01, ^###^p<0.001 by one sample t test with a hypothetical value of zero. For each group, n≥7 mice in at least two independent infections.

## DISCUSSION

During bacteremia, *K. pneumoniae* virulence and fitness factors may act during (1) initial site invasion, (2) dissemination, and (3) bloodstream survival (3). Based on data from multiple infection models, we identified GmhB as important in the third phase of bacteremia: bloodstream survival. In a model of bacteremic pneumonia, GmhB was dispensable for lung fitness but critical for fitness in the spleen. In *ex vivo* growth assays, GmhB was specifically important for spleen fitness. Furthermore, GmhB was also required by *E. coli* and *C. freundii* for bloodstream survival. Overall, this study indicates that GmhB is a conserved Gram-negative bloodstream survival factor.

Distinguishing the three pathogenesis phases of Gram-negative bacteremia can be difficult using *in vivo* infection models. While bacteremic pneumonia modeling indicated a role for GmhB in the latter two phases of bacteremia (Figure 1A), dissemination and bloodstream survival are difficult to separate experimentally since these processes occur simultaneously. To probe late phases individually, a dissemination independent model of direct bacteremia was utilized and confirmed a role for GmhB during bloodstream survival (Figure 1F). However, we cannot rule out a specific role in dissemination. Indeed, the greater Δ*gmhB* fitness defect observed in spleen and blood during bacteremic pneumonia compared to direct bacteremia suggests a role for GmhB in both dissemination and survival (Figure 1A, F). Lung dissemination mechanisms for *Pseudomonas aeruginosa* have been described and rely on exotoxins and the type 3 secretion system for killing host cells to gain bloodstream access (44–46). *K. pneumoniae* does not encode these factors (47). Instead, lung dissemination in *Klebsiella* requires a different host-pathogen interaction, where *K. pneumoniae* siderophores activate epithelial HIF-1α that is in turn required for dissemination (12). The precise mechanism of, and additional factors required for, dissemination from the lung is unclear.

GmhB is involved in the biosynthesis of ADP-heptose, a metabolite detected in host cytosol that initiates inflammation through the ALPK1/TIFA/NF-κB axis (28–31, 48, 49). GmhB dephosphorylates HBP to yield HMP1, which is converted into ADP-heptose. In the present study, GmhB was dispensable for normal inflammation during pneumonia as determined by immune cell recruitment and cytokines signatures associated with ALPK1/TIFA/NF-κB signaling. Therefore, lung inflammation elicited by *K. pneumoniae* may not require ADP-heptose or may be activated by other *K. pneumoniae* PAMPs. The minor differences in the LPS electrophoresis pattern in the absence of GmhB indicates that, as in *E. coli* (25, 31), *K. pneumoniae* possesses an unknown mechanism with partially redundant GmhB function (Figure 5). In the absence of GmhB, this mechanism may produce sufficient ADP-heptose to induce inflammation via the ALPK1/TIFA/NF-κB axis, leading to normal inflammation observed in Figure 2.

*K. pneumoniae* LPS O-antigen is required for serum resistance (14), but its role in lung fitness may vary. The strain KPPR1 requires LPS O-antigen for initial site lung fitness, while it is dispensable for the strain 5215R (13, 50). In *Salmonella* Typhimurium, complete abrogation of ADP-heptose integration into LPS results in a molecule lacking core and O-antigen (17, 18) and displays a rough phenotype. Here, GmhB was required for normal LPS biosynthesis but was not absolutely required for production of full length LPS containing O-antigen. Additionally, KPPR1 retained high levels of hypermucoviscosity in the absence of GmhB. Therefore, GmhB appears to maximize ADP-heptose biosynthesis and contribute to wild-type levels of LPS inner core production. Future work should discern how individual components of the LPS molecule contribute to bloodstream fitness and pathogenicity.

GmhB may be crucial under conditions where rapid LPS production is necessary. During murine bacteremia, *K. pneumoniae* exhibits exponential replication in the spleen at 24 hours (51). Rapid replication requires substantial LPS export and, in the absence of GmhB, lower abundance of normal LPS may be produced. This may leave Gram-negative species more susceptible to killing by host defenses, such as phagocytosis by immune cells. Our data supports differential requirements of capsule and LPS in site-specific fitness. The requirement of GmhB for fitness in the spleen *in vivo* and *ex vivo*, but dispensability for human serum resistance and lung and liver fitness *in vivo* and *ex vivo*, indicates that site specific immune cells like splenic macrophages may be required for *K. pneumoniae* clearance during bacteremia. In contrast, RfaH, necessary for capsule production and hypermucoviscocity, is dispensable for *ex vivo* spleen, liver and lung fitness but required for human serum resistance and *in vivo* lung fitness (11). This suggests that there are distinct interactions between *Klebsiella* and host defenses at each site of infection that require different *Klebsiella* virulence factors.

This study is limited by the validation rate of the InSeq selection process. Each InSeq model requires consideration of experimental bottlenecks to assess the maximum transposon library complexity which can be utilized (52, 53). Since Lcn2 restricts *K. pneumoniae* to the pulmonary space (23), *Lcn2*^-/-^ mice were used to relax the bottleneck between the lung and spleen, accommodating use of a complex *K. pneumoniae* transposon library that increased the number of disrupted genes. However, only one of the six hits chosen for validation significantly impacted bacteremia pathogenesis, suggesting that stochastic loss from a bottleneck still generated a high rate of false positive hits. The gene *prlC*, which in validation studies was an initial site fitness factor, encodes an oligopeptidase that may be important during lung infection. In future studies, this bottleneck could be mitigated by splitting the transposon library into smaller pools and increasing the number of replicates for each pool.

Based on InSeq studies and validation with isogenic mutants, GmhB is a conserved fitness factor across multiple species that cause bacteremia. Here, we confirmed a role for GmhB in bloodstream fitness for *K. pneumoniae*, *E. coli*, and *C. freundii*. InSeq analysis of *C. freundii* bacteremia fitness factors also indicated a role for GmhB in bloodstream fitness (56). Whereas GmhB is conditionally essential in these species, in *S. marcescens*, GmhB appears to be essential for growth (43). This consistent requirement for bloodstream survival makes GmhB and core LPS synthesis pathways attractive candidates for novel therapeutics to treat bacteremia.

## MATERIALS AND METHODS

### Transposon insertion site sequencing analysis (InSeq)

Construction of the *K. pneumoniae* transposon library using the pSAM_Cam plasmid and InSeq analysis was described previously (11). Briefly, after infection with the *K. pneumoniae* transposon library, CFU from total organ homogenate were recovered. DNA from recovered transposon mutants was extracted and fragments were prepared for Illumina sequencing using previously detailed methods (57). All transposon sequencing files are available from the NCBI SRA database (https://www.ncbi.nlm.nih.gov/sra, PRJNA270801).

### Bacterial strains and media

Reagents were sourced from Sigma-Aldrich (St. Louis, MO) unless otherwise noted. *K. pneumoniae* strains were cultured overnight in Luria-Bertani (LB, Fisher Bioreagents, Ottawa, ON) broth at 37°C shaking or grown on LB agar (Fisher Bioreagents) plates at 30°C. *E. coli* CFT073 (58) and *C. freundii* UMH14 (56) strains were cultured overnight in LB broth shaking or grown on LB agar plates at 37°C. Media for isogenic knockout strains and transposon mutants was supplemented with 40μg/mL kanamycin and pACYC was selected with 50μg/mL chloramphenicol.

Isogenic knockouts were constructed using Lambda Red mutagenesis and electrocompetent KPPR1 as previously described (11, 22). In short, electrocompetent *K. pneumoniae* carrying the pKD46 plasmid was prepared by an overnight culture at 30°C and diluted the following day 1:50 in LB broth containing 50μg/mL spectinomycin, 50mM L-arabinose, 0.5mM EDTA (Promega, Madison, WI), and 10μM salicyclic acid until reaching exponential phase, defined by an OD_600_ of 0.5-0.6. Bacterial cells were cooled on ice for 30 minutes, followed by centrifugation at 8,000xg for 15 minutes at 4°C. Pellets were washed serially with 50mL of 1mM HEPES pH 7.4 (Gibco, Grand Island, NY), 50mL diH_2_O, and 20mL 10% glycerol before making a final resuspension at 2-3×10^10^ in 10% glycerol. To generate gene-specific target site fragments for Lambda Red mutagenesis, a kanamycin resistance cassette was amplified from the pKD4 plasmid with primers also containing 65 base pair regions of homology to the chromosome flanking the *gmhB* open reading frame. The fragment was electroporated into competent KPPR1 containing pKD46 plasmid and transformants were selected on LB agar containing kanamycin after overnight incubation at 37°C. All KPPR1 isogenic knockouts were confirmed by colony PCR using gene internal and flanking primers. The *C. freundii* UMH14:Δ*gmhB* strain was constructed using Lambda Red mutagenesis as follows: Electrocompetent *C. freundii* UMH14 maintaining the pSIM18 recombination plasmid were prepared by harvesting exponentially growing cells cultured in YENB media supplemented with 200 μg/mL hygromycin grown at 30°C with aeration. To induce expression of pSIM18, the temperature was shifted to 42°C for 20 minutes and then the culture pelleted at 5,000xg for 10 minutes at 4°C. Cells were washed twice in cold 10% glycerol and resuspended in 100μL cold 10% glycerol before storage at −80°C. A gene-specific kanamycin resistance cassette was amplified from the pKD4 plasmid using primers containing 40 base pair regions of homology to the chromosome flanking the UMH14 *gmhB* open reading frame. This fragment was electroporated into UMH14 pSIM18 electrocompetent cells which were then recovered in LB media for 1 hour at 37°C and plated on LB agar containing kanamycin and incubated at 37°C overnight. UMH14:Δ*gmhB* was confirmed by Sanger sequencing and curing of the pSIM18 recombineering plasmid was confirmed by a restoration of hygromycin sensitivity. The primers used in this study are detailed in Supplementary Table 2.

The KPPR1 *gmhB* complementation plasmid, pACYC_*gmhB*_, was generated by two fragment Gibson assembly using NEBuilder HiFi DNA Assembly Master Mix (New England Biolabs, Ipswich, MA). The plasmid pACYC184 (pACYC_ev_; empty vector) was linearized by BamHI and HindIII (New England Biolabs). The *gmhB* locus, including a 500 bp region upstream of the open reading frame was amplified by PCR from KPPR1 (GCF_000755605.1, nucleotides 2,380,173 – 2,379,086) with primers containing homology to linearized pACYCev, described above. The plasmid and *gmhB* containing PCR product were mixed in a 1:2 ratio and Gibson assembly was performed following the manufacture’s protocol. The resulting Gibson product was electroporated and maintained in *E. coli* TOP10 cells (New England Biolabs) and the final construct (pACYC_*gmhB*_) was confirmed using Sanger sequencing. *pACYC_gmh_B* and pACYC_ev_ were mobilized into KPPR1 and Δ*gmhB* by electroporation and plasmids were maintained in the presence 50μg/mL chloramphenicol.

### Murine bacteremia models

This study was performed using six-to ten-week old C57BL/6 mice (Jackson Laboratory, Bar Harbor, ME) with careful adherence to humane animal handling recommendations (59) and the study was approved by the University of Michigan Institutional Animal Care and Use Committee (protocol: PRO00009406). As a model of bacteremic pneumonia, mice were anesthetized with isoflurane and 1×10^6^ CFU *K. pneumoniae* in a 50μL volume was administered retropharyngeally. For intraperitoneal bacteremia, mice were injected with 1×10^3^ CFU *K. pneumoniae* in a 100μL volume administered to the peritoneal cavity. For direct bacteremia, mice were injected with 1×10^5^ CFU *K. pneumoniae* in a 100μL volume administered via tail vein injection (60). For all models, overnight LB cultures of *K. pneumoniae* were centrifuged, resuspended, and adjusted to the proper concentration in PBS. Twenty-four hours post infection, mice were euthanized by carbon dioxide asphyxiation prior to collection of blood, lung, spleen, liver, or peritoneal fluid. Whole blood was collected by cardiac puncture and dispensed into heparin coated tubes (BD, Franklin Lakes, NJ). Peritoneal fluid was collected by dispensing 3mL PBS into the peritoneal cavity followed by recollection. After collection, all organs were homogenized in PBS. To determine bacterial density, all sites were serially diluted and CFU measured by quantitative plating on LB agar with appropriate antibiotics. To calculate competitive indices, mice were infected with a 1:1 ratio of *K. pneumoniae* wild-type KPPR1 or isogenic mutant strains. Total CFU were determined by LB agar quantitative plating and mutant strain CFU were quantified by plating on LB agar with appropriate antibiotics. The competitive index was defined as CFU from: (mutant output/wild-type output)/(mutant input/wild-type input).

To model *E. coli* bacteremia, mice were inoculated with a 1:1 mixture of CFT073:tn::*gmhB* for a total of 1×10^7^ CFU in a 100μL volume administered via tail vein injection. To model *C. freundii* bacteremia, UMH14 and UMH14:Δ*gmhB* stationary phase cultures were back diluted (1:100) into fresh LB media and grown to late exponential phase at 37° C with aeration. These cultures were centrifuged at 5,000xg for 10 minutes at 4°C, and the pellets were suspended in cold PBS to 5×10^8^ CFU/mL for UMH14 and 1×10^9^ CFU/mL for UMH14:Δ*gmhB* and then combined 1:1. 100μL of the combined suspension, which constituted a total inoculum of 7.5×10^7^ CFU at a 1:2 CFU ratio of wild-type to mutant, was administered by tail vein injection. For *E. coli* and *C. freundii*, enumeration of total CFU per organ was performed with serial dilution plating as above (using 50μg/mL kanamycin for *C. freundii*), and the calculation of competitive indices were determined as described above.

### Flow cytometry

Lung homogenate was collected 24-hours post infection with either KPPR1 or Δ*gmhB* in the bacteremic pneumonia model. Lungs were prepared for flow cytometry using single cell suspensions as previously described (61). In short, lungs were resected, minced, and digested in a buffer containing complete DMEM (10% FBS), 15mg/mL collagenase A (Roche, Basel, Switzerland) and 2000 units of DNase for 30 minutes at 37°C. Following digestion, samples were disrupted by repeated aspiration through a 10mL syringe. Leukocytes were isolated by centrifuging disrupted tissue through a 20% Percoll Solution (2,000xg for 20 minutes). 1.5 x 10^6^ leukocytes were stained with diluted antibody for 30 minutes on ice before analysis on a BD Fortessa Cytometer. Staining antibodies included: BV650-CD11b (clone M1/70), BV421-I-Ab (MHCII clone AF6-120.1), APC-Cy7-SiglecF (clone E50-2440), purchased form BD Horizon; PE-eFluor610-CD11c (clone N418), purchased from eBioscience; BV605-CD62L (clone MEL-14), BV510-Cx3CR1 (clone SA011F11), AlexaFluor700-CD45 (clone I3/2.3), PE-CD64 (clone X54-5/7.1), PerCP-Cy5.5-CD24 (clone M1/69), PE-Cy7-Ly6C (clone HK1.4), BV570-Ly6G (clone 1A8), APC-CD115 (clone AFS98), purchased from Biolegend. Visualization of cell populations was assembled using FlowJo (Version 10.7.2).

### Cytokine ELISAs

Mice were infected with either KPPR1 or Δ*gmhB* using the bacteremic pneumonia model and lungs were homogenized with tissue protein extraction reagent (T-PER, Fisher). Homogenate was centrifuged at 500xg for 5 minutes and the supernatant was analyzed for cytokine abundance by the University of Michigan Rogel Cancer Center Immunology Core Facility using enzyme-linked immunosorbent assay (ELISA).

### Serum killing and growth assays

To measure serum susceptibility, 1×10^5^ CFU of stationary phase *K. pneumoniae* was added to 100% active human (Invitrogen, Waltham, MA) or C57B/L6 murine serum (Invitrogen). Plates were incubated at 37°C for three hours, and killing was measured by serial dilutions and quantitative culture at t=0 and t=3. To assess growth, overnight LB broth *K. pneumoniae* cultures were adjusted to 1×10^7^ CF/mL in M9 salts plus 20% human serum in a 96-well dish. Samples were incubated at 37°C and OD_600_ readings were measured every 15 minutes using an Eon microplate reader and Gen5 software (Version 2.0, BioTek, Winooski, VT).

### *Ex vivo* survival assay

Spleen, liver, and lung from uninfected mice were homogenized in 2mL PBS. Overnight LB broth *K. pneumoniae* cultures were adjusted to 1×10^6^ CFU/mL in PBS and mixed 1:1 for competitive growth. From the bacterial suspension, 10μL was added to 90μL of organ homogenate for a final concentration of 1×10^5^ CFU/mL and incubated for 3 hours at 37°C. Survival was measured by serial dilutions and quantitative culture at t=0 and t=3.

### LPS isolation and electrophoresis

LPS from 1×10^9^ CFU of each strain of interest was isolated using the Sigma Lipopolysaccharide Isolation Kit according to the manufacturer’s instructions. Electrophoresis was performed using a 4-20% mini-PROTEAN TGX Precast gel (Bio-Rad, Hercules, CA). LPS was visualized by staining with the Pro-Q Emerald 300 Lipopolysaccharide Gel Stain Kit (Molecular Probes, Eugene, OR).

### Statistical analysis

Each *in vivo* experiment was performed in at least two independent infections, and each *in vitro* experiment was an independent biological replicate. For each study, statistical significance was defined as a p-value <0.05 (GraphPad Software, LaJolla, CA) as determined by: one-sample test to assess differences from a hypothetical competitive index of zero, unpaired t test to assess differences between two groups, or ANOVA followed by Tukey’s multiple comparisons post-hoc test to assess differences among multiple groups.

## ACKNOWLEDGEMENTS

CLH is supported by the Lung Immunopathology Training Grant (T32HL007517); SJG is supported by R35HL144481; LVU is supported by the Molecular Mechanisms in Microbial Pathogenesis Training Program (32AI007528-21A1); GSB, HLTM, and MAB are supported by AI134731 from the National Institutes of Health.

The authors thank Mark T. Anderson for technical support in LPS isolation and electrophoresis. All authors disclose no conflicts of interest.

## SUPPLEMENTARY TABLES

**Supplemental Table 1.**
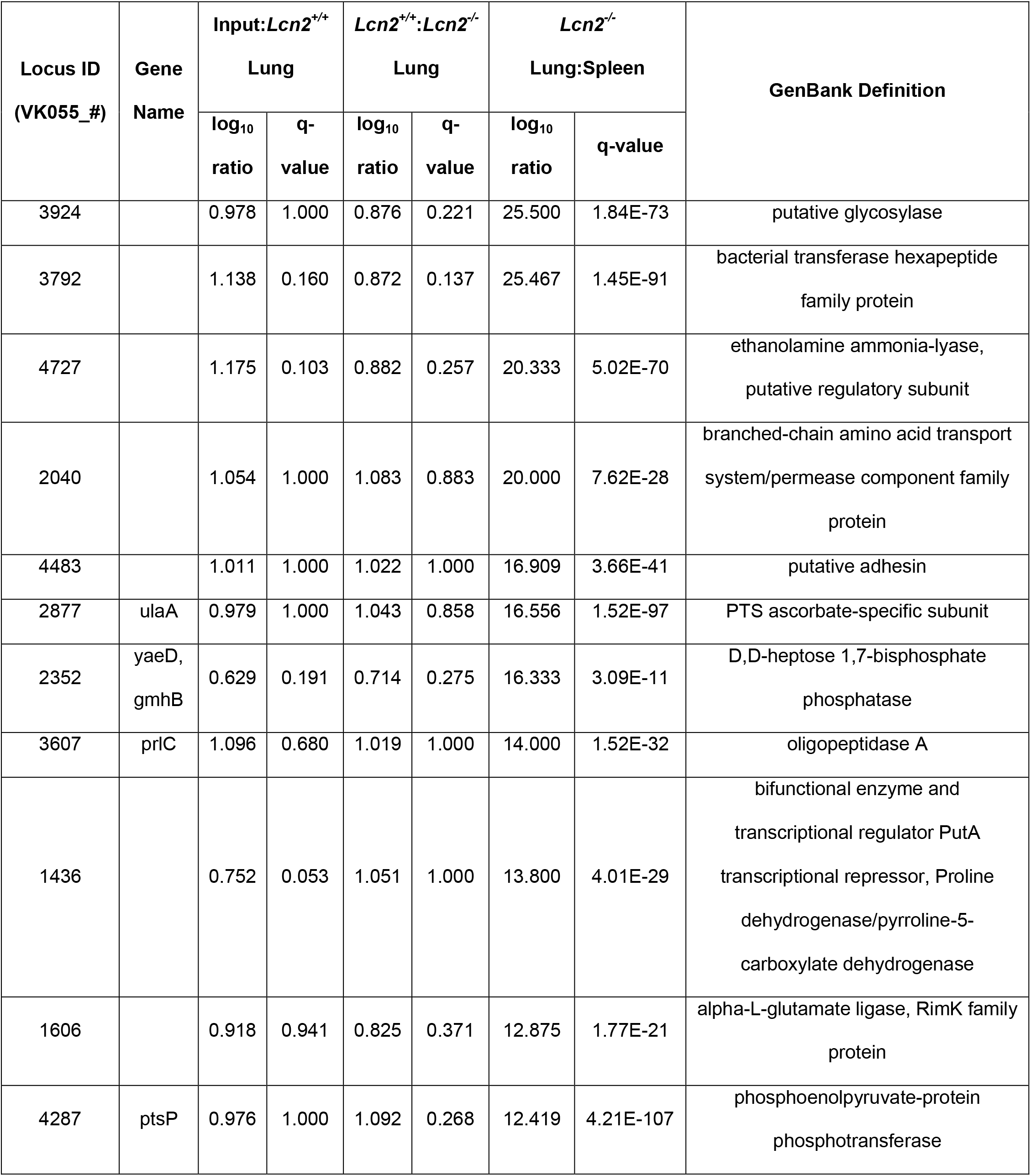
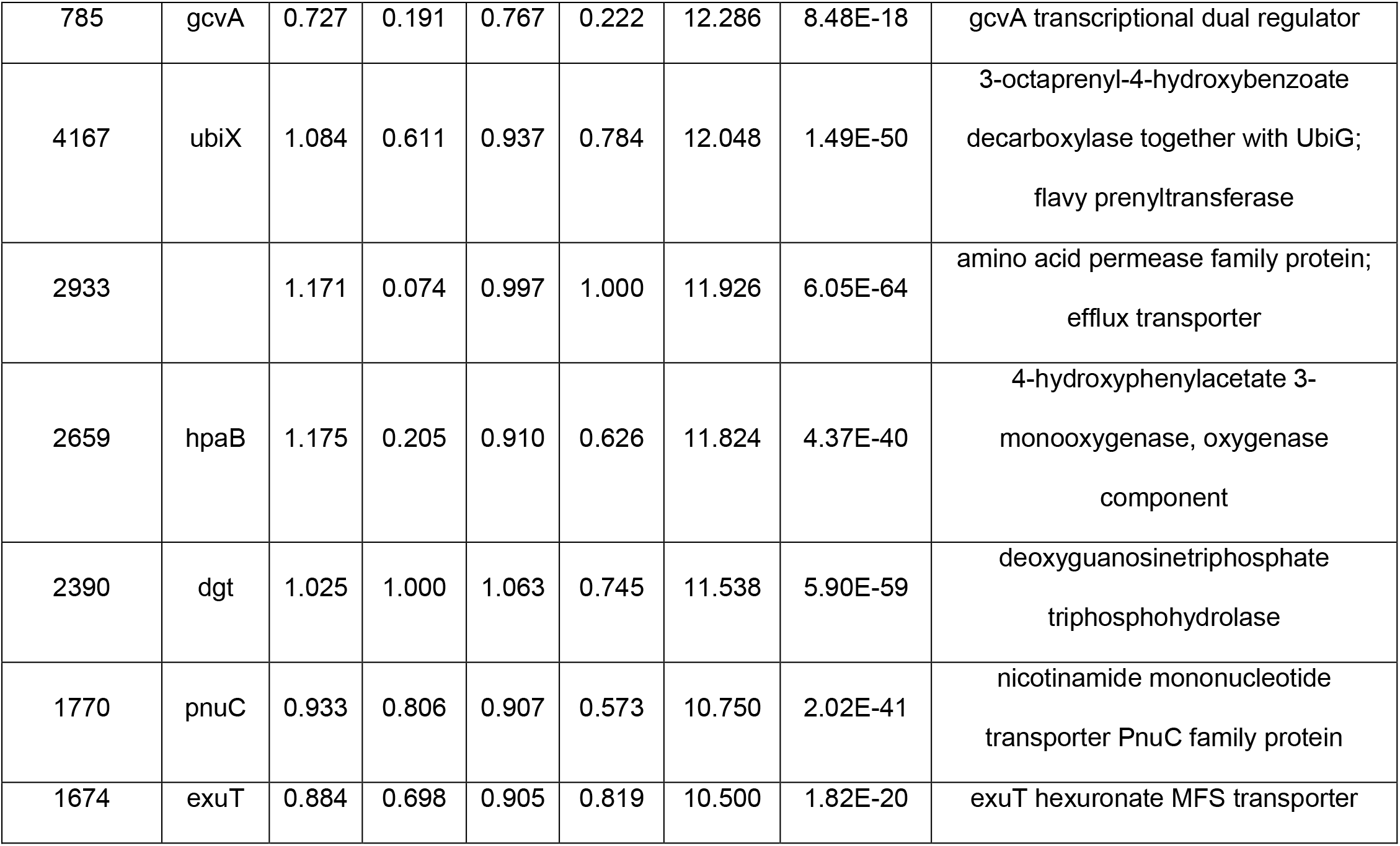
Factors identified by transposon insertion site sequencing (InSeq) for involvement in late phases of *K. pneumoniae* bacteremia.

## SUPPLEMENTARY FIGURES

**Supplementary Figure 1:**
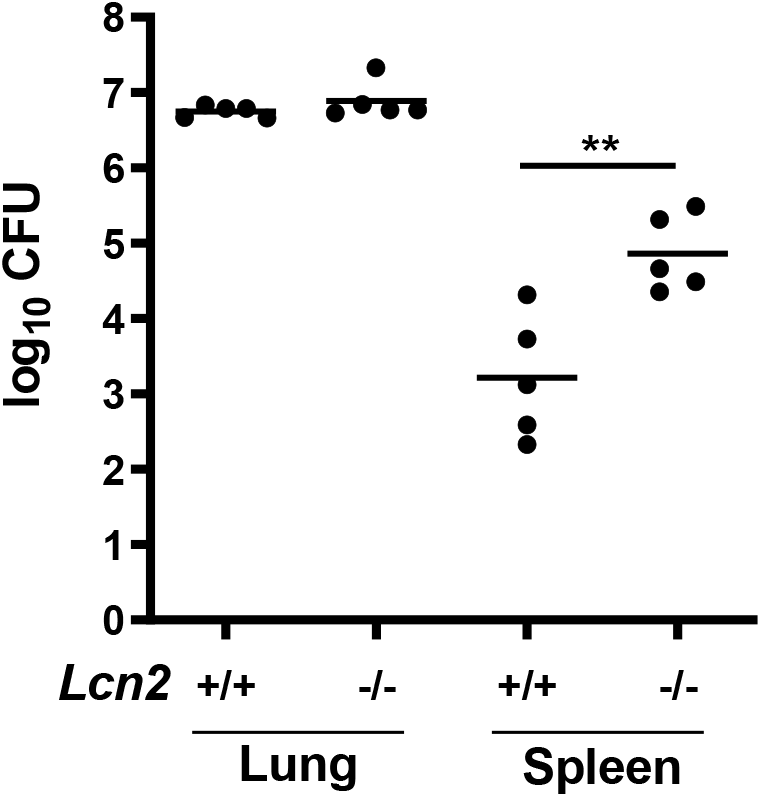
Lipocalin 2 restricts *K. pneumoniae* lung dissemination. To model pneumonia, 1×10^6^ CFU of a library of *K. pneumoniae* transposon mutants was administered retropharyngeally to *Lcn2*^+/+^ or *Lcn2*^-/-^ mice as previously reported (11). Mean log_10_ CFU is displayed for each organ at 24 hours post infection. **p<0.001 by unpaired t test. For each group, n=5 mice.

**Supplementary Figure 2.**
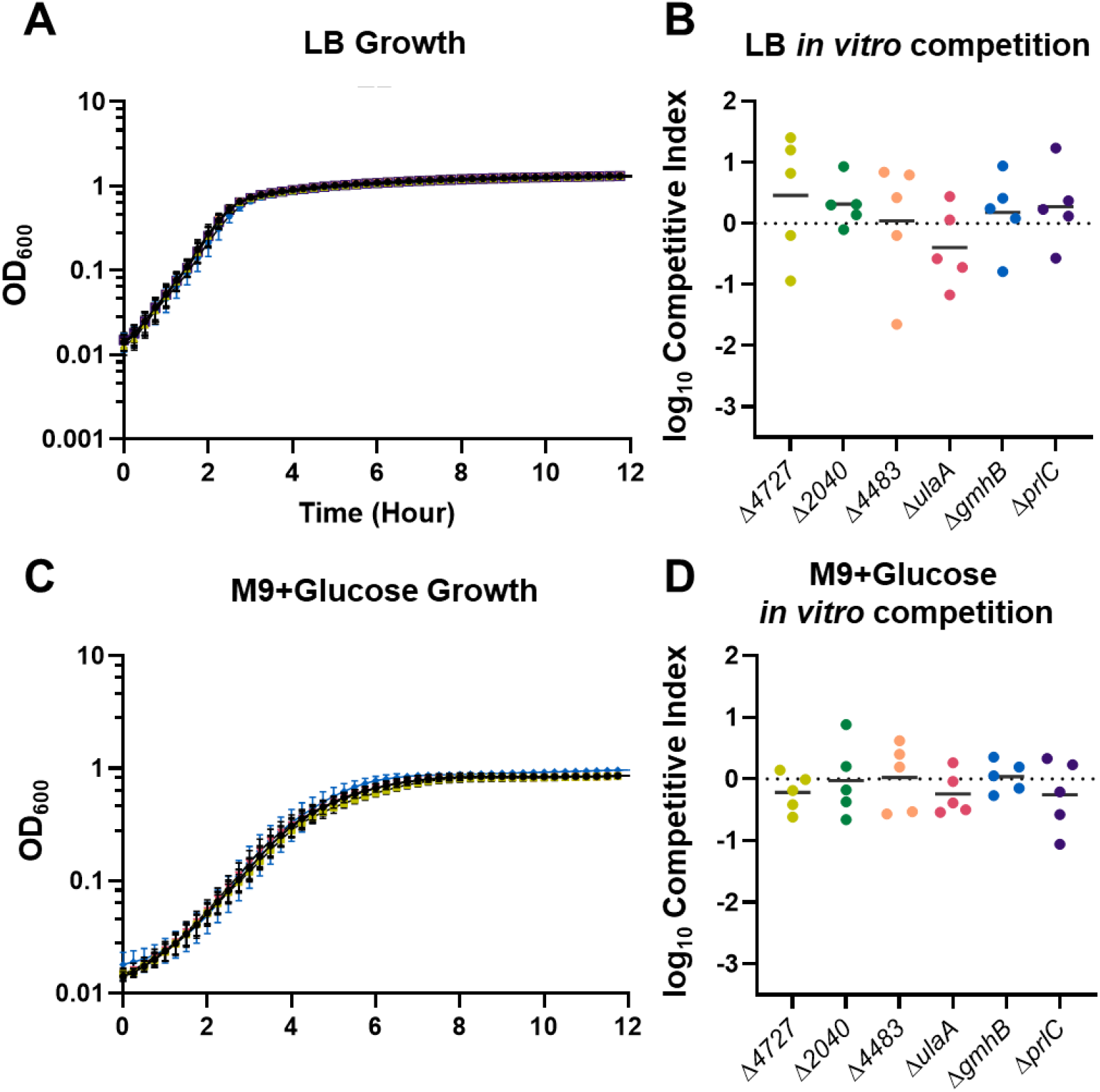
GmhB and other factors of interest are not required for *K. pneumoniae* replication *in vitro*. KPPR1 or isogenic knockouts were inoculated to a starting concentration of 1×10^7^ CFU/mL and monitored by optical density (OD_600_) in LB (A) and M9 with 0.9% glucose (M9+Glucose; C). KPPR1 and each mutant were combined 1:1 at a concentration of 1×10^6^ CFU/mL and incubated in LB (B) or M9+Glucose (D) and mean log_10_ competitive index compared to wild-type at 24 hours post inoculation is displayed (n=5). One-way ANOVA indicated no significant difference between strains for area under the curve after growth and one-sample t tests with a hypothetical value of zero showed no defect in competitive indices; for A and C, lines colors correspond to strains in B and D.

**Supplementary Figure 3.**
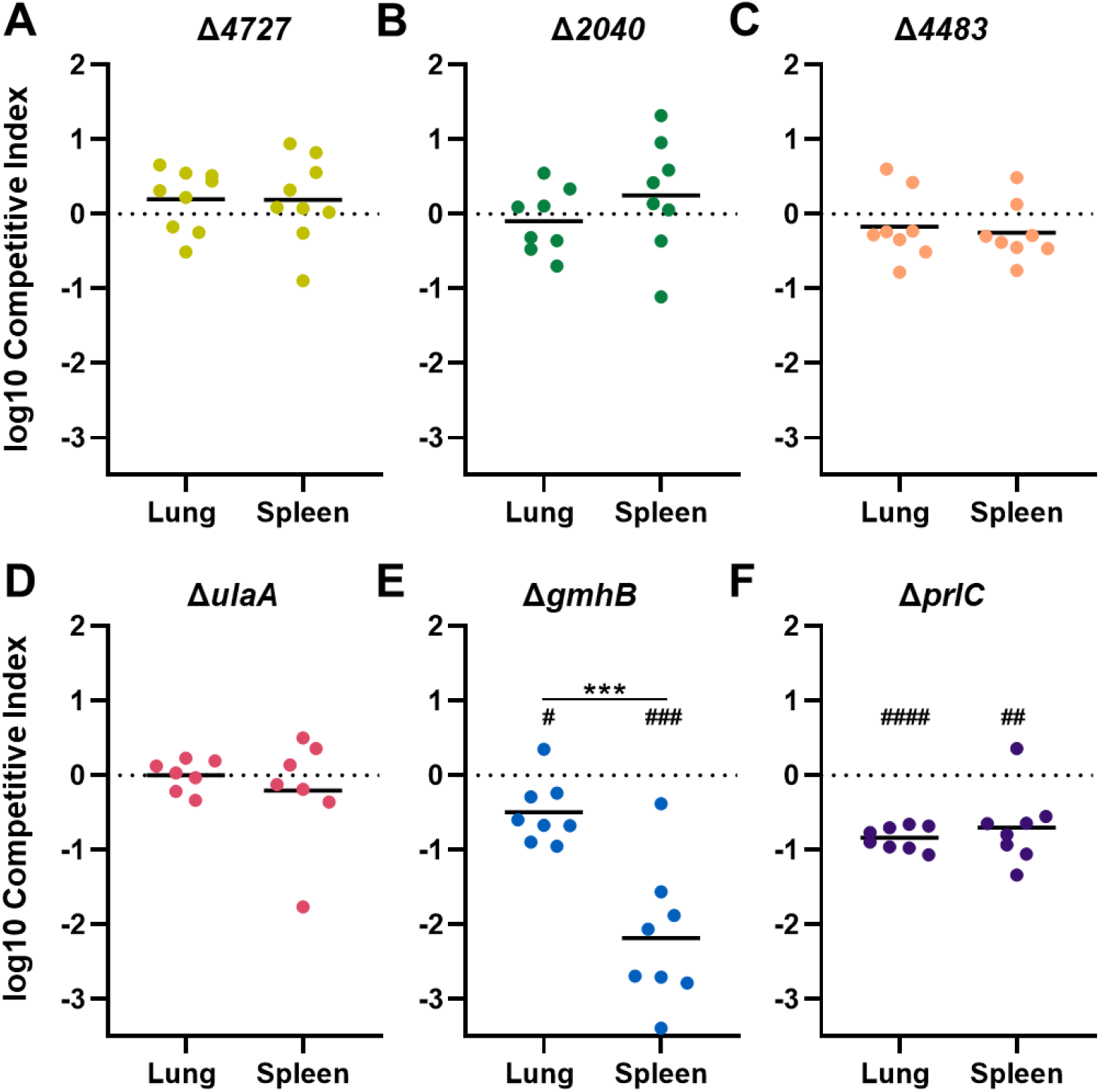
InSeq analysis reveals *K. pneumoniae* GmhB as enhancing late bacteremia fitness. Isogenic knockouts were constructed to validate the InSeq selection approach identifying dissemination and bloodstream survival factors (A-F). Each knockout was mixed 1:1 with KPPR1 for a final inoculum of 1×10^6^ CFU and administered in the pharynx of *Lcn2*^-/-^ mice. Mean log_10_ competitive index compared to wild-type at 24 hours post infection is displayed. ***p<0.001 by unpaired t test; ^#^p<0.05, ^##^p<0.01, ^###^p<0.001, ^####^p<0.0001 by one sample t test with a hypothetical value of zero. All statistical tests were performed on log-transformed data. For each group, n≥7 mice across at least two independent infections.

**Supplementary Figure 4.**
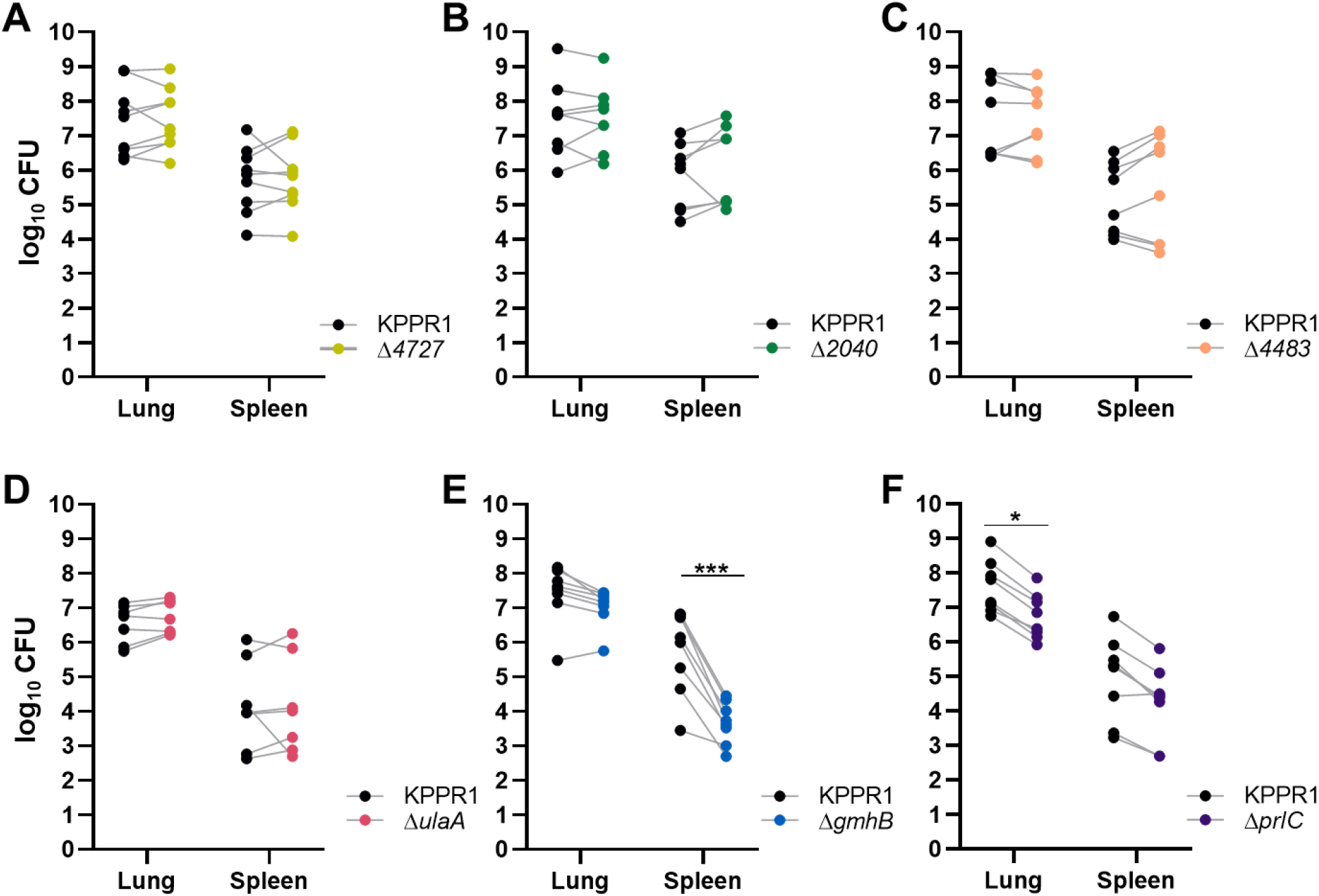
Bacterial burden summary for *in vivo* validation of transposon insertion site sequencing (InSeq). CFU per organ from mice 24 hours after inoculation with 1:1 mixture of isogenic knockouts and KPPR1 (1×10^6^ CFU total) administered in the pharynx of *Lcn2*^-/-^ mice is shown (A-F), corresponding to the competitive indices in Figure S3. *p<0.05, ***p<0.001 by unpaired t test. For each group, n≥7 mice in at least two independent infections.

**Supplementary Figure 5.**
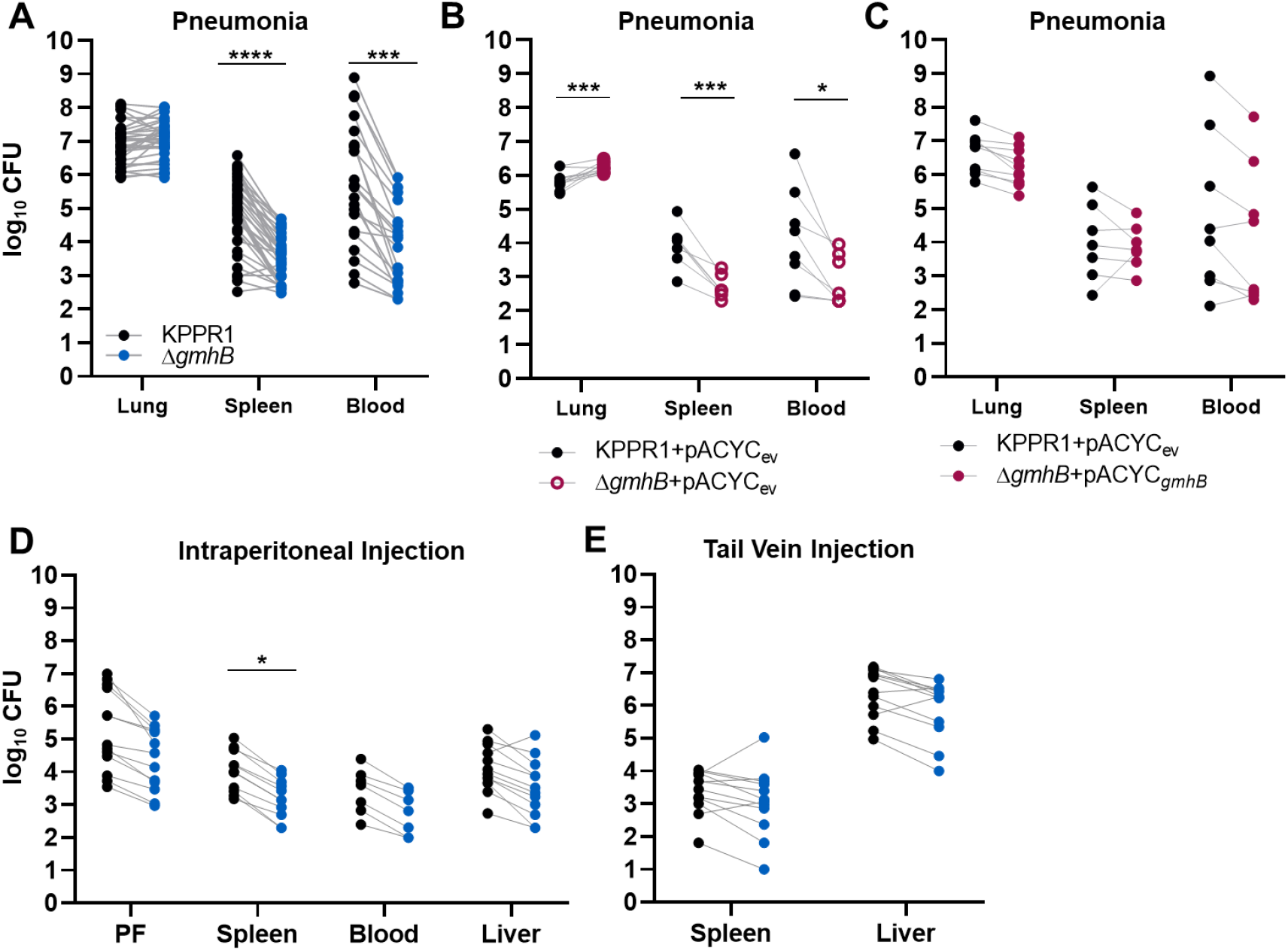
Bacterial burden summary for models of murine bacteremia. In a model of bacteremic pneumonia, mice were retropharyngeally inoculated with 1×10^6^ CFU *K. pnuemoniae* (A-C). To initiate dissemination from a lung-independent site, 1×10^3^ CFU was administered to the intraperitoneal cavity (D). For modeling direct bacteremia requiring no dissemination, 1×10^5^ CFU was administered via tail vein injection (E). The 1:1 inoculum consisted of KPPR1:Δ*gmhB* (A, D, E), KPPR1:Δ*gmhB* carrying empty pACYC vector (ev; B), or KPPR1_ev_:Δ*gmhB* with *gmhB* complementation provided on pACYC under control of the native *gmhB* promoter (Δ*gmhB*+pACYC_*gmhB*_; C). Log10 CFU burden for each site at 24 hours post infection is displayed, corresponding to competitive indices in Figure 1. *p<0.05, **p<0.01, ***p<0.001, ****p<0.0001 by unpaired t test. For each group, n≥7 mice in at least two independent infections.

**Supplemental Figure 6.**
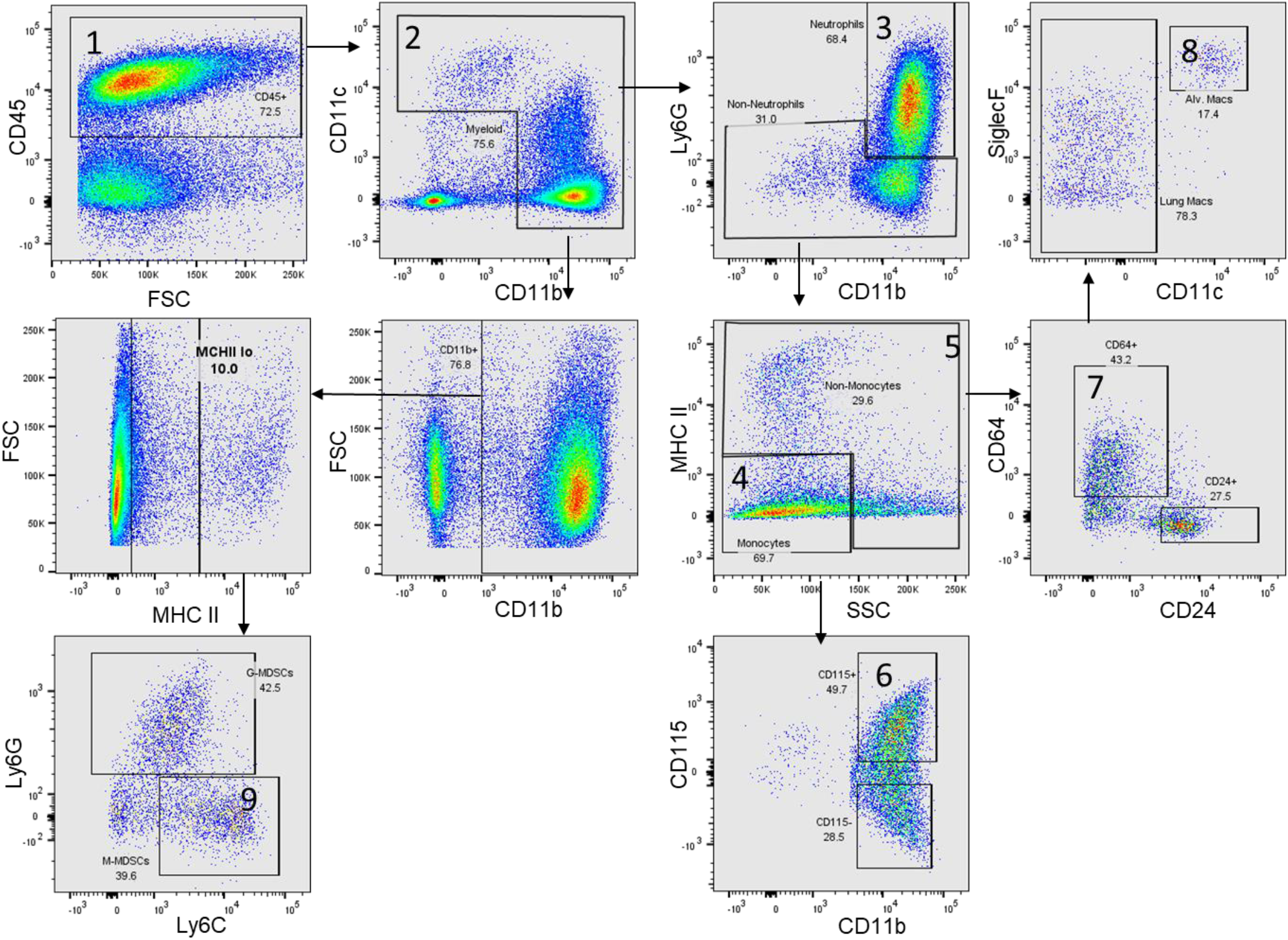
Gating Scheme for flow cytometry experiments. Single cell suspensions were generated from collagenase digested lungs as described. Following this, cell viability was assessed via trypan blue exclusion and was >90% for all samples. Cells were subsequently gated as follows: CD45^+^ (Gate 1), myeloid lineage cells: CD11b/c^+^ (Gate 2), neutrophils: Ly6G^+^ (Gate 3), putative monocytes: MHCII^-^, SSC^lo^ (Gate 4) or macrophage and DCs: MHCII^+/-^ SSC^hi^ (Gate 5), CD115^+^ Monocytes (Gate 6), macrophages: CD64^+^, CD24^-^ (Gate 7), alveolar macrophages: SiglecF^+^, CD11c^+^ (Gate 8). M-MDSCs: CD11b^+^, MCH^lo^, Ly6G^-^, Ly6C^+^ (Gate 9).

**Supplementary Figure 7.**
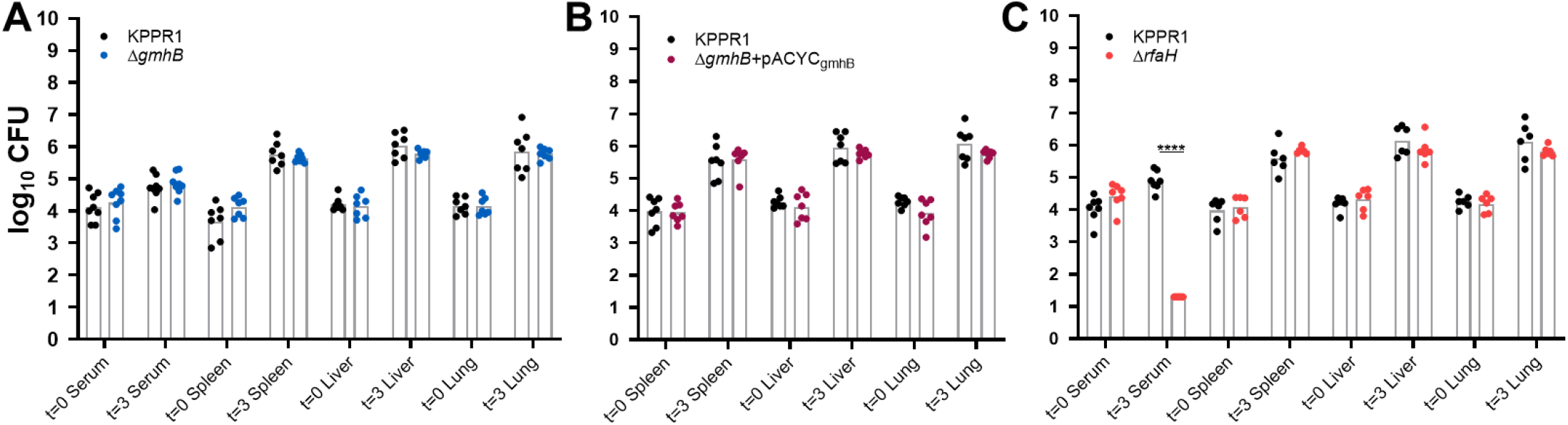
Bacterial counts from *ex vivo* killing assays. *Ex vivo* competition assays were performed in human serum and uninfected murine spleen, liver, or lung homogenate using 1:1 mixture of KPPR1 and either Δ*gmhB* (A), Δ*gmhB*+pACYC_*gmhB*_ (B), or Δ*rfaH* (C). Log10 recovered CFU following 0 hours (t=0) and three hours (t=3) of incubation in specified condition is displayed, corresponding to competitive indices in Figures 3 and 4. ****p<0.0001 by unpaired t test, for each group, n≥7.

**Supplementary Figure 8.**
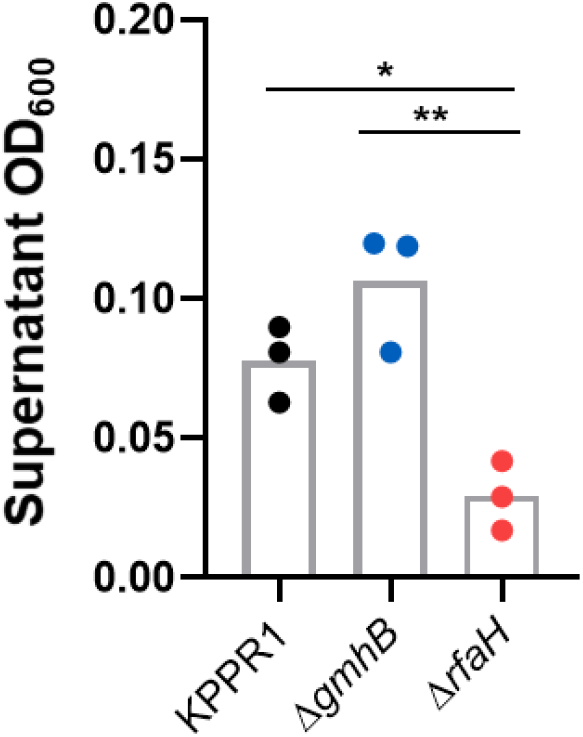
GmhB does not alter hypermucoviscosity. To assess hypermucoviscosity, overnight cultures were pelleted at 5,000xg for 15 minutes and adjusted to an OD_600_=1 in 1mL PBS. Normalized PBS suspensions were subsequently centrifuged at 1,000xg for 5 minutes and the OD_600_ of the upper 900μL of supernatant was measured from three biological replicates.

**Supplementary Figure 9.**
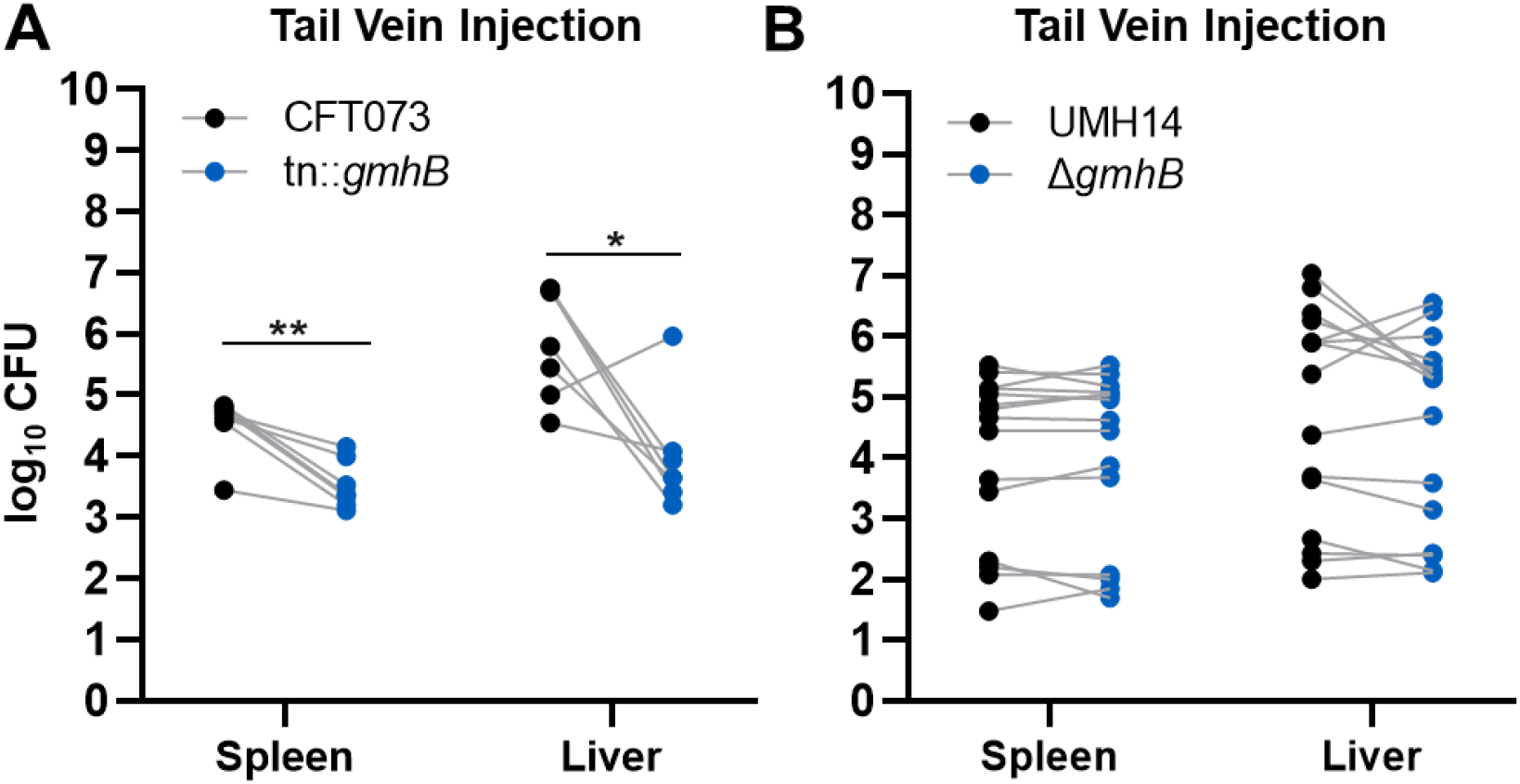
Bacterial burden summary for direct bacteremia with *E. coli* and *C. freundii*. In a model of direct bacteremia, 1×10^7^ CFU of *E. coli* CFT073 mixed 1:1 with CFT073:tn::*gmhB* (A) or *C. freundii* UMH14 mixed 1:2 with UMH14:Δ*gmhB* (B) was administered via tail vein injection. Log10 CFU burden for each site at 24 hours post infection is displayed, corresponding to competitive indices in Figure 6. *p<0.05, **p<0.01 by unpaired t test. For each group, n≥7 mice in at least two independent infections.

